# Temporal regulation of AgRP neurons mediates context-induced feeding

**DOI:** 10.1101/2023.01.30.526347

**Authors:** Felicia Reed, Harry Dempsey, Rachel E Clarke, Alex Reichenbach, Mathieu Mequinion, Romana Stark, Sasha Rawlinson, Claire J Foldi, Sarah H. Lockie, Zane B. Andrews

## Abstract

An environment can have a powerful influence over appetite and feeding behaviour. For example, an environmental context, which reliably predicts food, will increase the appetitive food drive to the same environment context. Interestingly, mice are required to be hungry to develop such a context-induced feeding (CIF) response, suggesting the neural circuits sensitive to hunger play an important role to associate an internal energy state with a particular environment context. Hunger-sensing Agouti related peptide (AgRP) neurons are activated by circulating signals of energy deficit and reset to a silenced state by gut feedback mechanisms following food consumption. We hypothesised that AgRP neurons are both necessary and sufficient to drive CIF in the absence of hunger. While fasting increased CIF, chemogenetic inhibition of AgRP neurons during context acquisition prevented this effect. Intriguingly, chemogenetic activation of AgRP neurons during context acquisition did not increase CIF, suggesting precise temporal firing properties may be required. Indeed, photostimulation of AgRP neurons, only during context exposure (ON-OFF in context), increased CIF. Moreover, AgRP photostimulation prior to context exposure, coupled with the termination of photostimulation in the context in the absence of food consumption, was sufficient to drive a subsequent CIF. Our results suggest that AgRP neurons regulate the acquisition of CIF when the temporal firing properties are matched to context exposure. These results further highlight that acute AgRP inhibition is a salient neural event underscoring the effect of hunger on associative learning.

## Introduction

Hunger acts as a powerful incentive for an animal to interact resourcefully with its environment, and many higher-order cognitive processes likely evolved to support reliable food-finding under conditions of energy deficit. Hunger elicited by food restriction is frequently used as a motivational tool to accelerate acquisition of appetitive learning tasks in the laboratory (Beraldo et al., 2019; Heath et al., 2016; Horner et al., 2013; Ingram et al., 1987), despite limited knowledge of how hunger affects information processing in the brain. Indeed, hunger states influence the encoding of sensory information and improve memory function to both food and non-food related stimuli (Burgess et al., 2018; Critchley and Rolls, 1996; Radel and Clement-Guillotin, 2012; Shi et al., 2018; Sultson et al., 2019).

Hunger-associated conditions such as calorie restriction (CR), time-restricted feeding, or exogenous administration of ghrelin improve acquisition on simple object recognition tasks (Hornsby et al., 2015; Hsu et al., 2016; Pitsikas and Algeri, 1992), whereas conditions of diet-induced obesity or acute bouts of high calorie feeding impair performance on comparable tasks (McLean et al., 2018; Towers et al., 2020). Not only can hunger promote memory, but this link appears to be reciprocal; in that memories of past experiences influence current levels of appetite. For example, situations previously paired with hunger can elicit conditioned feeding responses in the absence of energy deficit (Bouton, 2011; Stern et al., 2020; Weingarten, 1983), while situations signalling aversive outcomes can suppress appetite and food intake in the face of energy deficit (Yang et al., 2021a). Despite the large number of studies demonstrating the effect of appetite on memory, the neurobiological processes and neural circuits linking appetite and memory remain poorly understood.

Hunger is associated with increased activity of AgRP neurons located in the arcuate nucleus of the hypothalamus (Arc) (Mandelblat-Cerf et al., 2015; Siemian et al., 2021; Wu et al., 2014). AgRP neurons co-express GABA, neuropeptide Y (NPY), and AgRP peptide; all of which contribute to feeding, albeit through different downstream targets and across varying timescales (Engstrom Ruud et al., 2020; Krashes et al., 2013). Chemogenetic and optogenetic activation of these neurons is sufficient to drive feeding and appetitive behaviours comparable to those elicited by fasting (Aponte et al., 2011; Krashes et al., 2011), and activation of these neurons in fed mice induces recall of a location-specific memory acquired under food restriction (Burnett et al., 2016) or a specific operant behavioural sequence previously rewarded under food restricted conditions (Siemian et al., 2021). In addition, AgRP neuronal activation influences mood (Dietrich et al., 2015; Fang et al., 2021; Li et al., 2019), dopamine driven motivation (Reichenbach et al., 2022), reward, reinforcement and valence attribution (Alhadeff et al., 2019; Betley et al., 2015; Chen et al., 2016), as well as associative learning (Berrios et al., 2021; Betley et al., 2015; Livneh et al., 2017).

Under physiological conditions, AgRP neurons are silenced prior to food consumption in response to learned sensory cues that predict food (Betley et al., 2015; Chen et al., 2015; Mandelblat-Cerf et al., 2015; Reichenbach et al., 2022). Moreover, the fall in AgRP activity depends on both the current energy need of the animal and the calorie content of available food, with the greatest suppression occurring in fasted mice or when presented palatable food or calorie-containing gels (Chen et al., 2015; Su et al., 2017). A sustained fall in AgRP activity requires both metabolic sensing in AgRP neurons (Reichenbach et al., 2022), as well as gastrointestinal feedback via increased gut peptide signalling and gut-brain neural feedback (Beutler et al., 2017; Goldstein et al., 2021; Su et al., 2017), which act to confirm calorie intake and content. For example, when mice are re-exposed to novel flavoured food gels, a sustained suppression of AgRP neuronal activity only occurs in response to caloric gels, and only when mice have been allowed to consume these gels – clearly demonstrating the necessity of nutrient dependent feedback in this process (Su et al., 2017). Moreover, inactivation of an afferent inhibitory GABAergic dorsomedial hypothalamic (DMH) pathway to AgRP neurons delays learning in a visual cue-initiated food-seeking task (Berrios et al., 2021). Thus, the acute inhibitory control of AgRP neurons seems important to guide learning. Based on these findings, we have recently argued that sensory and temporal integration of food or food cues regulate AgRP neurons in a predictive and experience-dependent manner (Reed et al., 2022). Therefore, memories of past feeding episodes, prompted by food-associated cues, may regulate AgRP activity in manner proportionate to expected outcomes and influence future feeding episodes. Collectively the evidence presented above suggests AgRP neurons provide a critical bidirectional link between hunger and memory. Given that AgRP neurons sense interoceptive hunger states (Siemian et al., 2021) and hunger elicits conditioned feeding responses paired with distinct contextual information (Bouton, 2011; Petrovich et al., 2007; Stern et al., 2020; Weingarten, 1983), we hypothesised that AgRP neurons encode interoceptive- and context-dependent information into conditioned feeding responses. Moreover, the rapid inhibition of AgRP neurons in response to the sensory detection of food suggests that the temporal silencing of AgRP neurons themselves is critical to facilitate the conditioning to environmental cues. Therefore, we assessed specific temporal role of AgRP neuronal manipulation with either chemogenetics or optogenetics in response to environmental cues using a context-induced feeding assay, independent of energy need.

## Methods

### Mice and housing

All experiments were conducted in accordance with the Monash Animal Ethics Committee guidelines, (MARP 18012). Mice were group housed under standard conditions (12:12 light-dark cycle, lights off at 7pm) and given *ad libitum* access to standard chow diet (no. 8720610, Barastoc Stockfeeds, Victoria, Australia) prior to surgical intervention.

*Agrp-ires-cre* mice on a C57BL/6J background (*Agrp*^*tm1(cre)Low/J*^) were obtained from Jackson Laboratory (stock no. 012899; The Jackson Laboratory, Maine, USA) and crossed with C57BL/6J mice obtained from the Monash Animal Research Platform (MARP; Clayton, AU). Male offspring heterozygous or wild-type (wt) for the cre allele were used for experiments (*Agrp*^cre/wt^ and *Agrp*^wt/wt^; respectively). Following surgeries, all mice were single housed and given free access to Irradiated Rat and Mouse Maintenance Chow (4.8 % fat, 19 % protein, 60 % carbohydrates; Specialty Feeds, Australia) for a minimum of 2 weeks prior to experimentation.

Adult male C57Bl/6J mice aged 12 weeks on arrival were obtained from MARP and group housed 2-4 mice per cage under the same standard conditions and allowed to acclimate to the behavioural facility for a minimum of 7 days before experimentation.

### Stereotaxic Surgery

Stereotaxic surgeries were performed on adult male *Agrp*^cre/wt^ and *Agrp*^wt/wt^ littermates at least 10 weeks of age. For all surgeries, *Agrp*^cre/wt^ and *Agrp*^wt/wt^ mice received bilateral injections of AAV (∼2.0 × 10^12^ vg/ml), 200nl/side infused at a rate of 40nl/min and allowed to rest for 5 minutes post-infusion. All viruses were purchased from Addgene (AAV5-hSyn-DIO-hM4D(Gi)-mCherry, #50475; AAV5-hSyn-DIO-hM3D(Gq)-mCherry, #50474; AAV9 hSynapsin-FLEX-soCoChR-GFP, #107712-AAV9) and injected into the mid-arcuate nucleus using a 2 *μ*l Hamilton Neuros Syringe (Stereotaxic co-ordinates, from bregma: AP: -1.6mm, ML: +/- 0.2mm, DV: -5.5/-5.6mm from brain surface). For optogenetic experiments, mice additionally received a unilateral fiber optic implant into the left hemisphere with 5.8 mm long ferrule capped fibers (400 µm core, NA 0.48, MF1.25 or MF 2.5 400/430-0.48; Doric studios, Canada) above the injection site and fixed in place with G-flow dental adhesive (GBond, GC Dental, Japan). Mice had a minimum of 2 weeks post-surgical recovery time, also permitting time for viral transfection.

### Drugs and compounds

Clozapine-N-oxide (CNO) (Carbosynth, UK) was dissolved in sterile saline (0.9 % NaCl) for injections. For Designer Receptor Exclusively Activated by Designer Drug (DREADD) activation studies (hM3Dq), a concentration of 1 mg/kg was used, whereas 3 mg/kg was used for inhibition studies (hM4Di). CNO was injected intraperitoneally (i.p) at a volume of 10 ul/g of body weight.

Purified rat ghrelin (R&D Systems, USA) was dissolved in saline on the day of injection. Experimental mice received a dose of 1 mg/kg, which significantly increased food intake, delivered i.p. at a volume of 10 ul/g of body weight.

### Optogenetics

Optogenetic experiments were conducted using optical components from Doric Lenses (Quebec, Canada). 10 mW of 465nm blue light was delivered from an LED driver to a connectorized LED coupled with a fibre optic rotary joint. Light output power was standarised to 10mW per setup using an Optical Power Meter (PM100D, ThorLabs, NJ USA) measured at the end of the fiber optic patch cable. For CIF and home cage feeding experiments, the following stimulation parameters were used: 20Hz pulses delivered every 1 in 4 seconds (1s ON, 3s OFF), based on the reported firing frequency of AgRP neurons during fasting *in vivo* (Mandelblat-Cerf et al., 2015). Pulse generation was controlled by an optogenetics TTL pulse generator using Doric Neuroscience Studio software.

### Functional validation of AgRP neuron activation with optogenetics or DREADDs

Chemogenetic or optogenetic activation of AgRP neurons drives robust food intake in fed mice (Aponte et al., 2011; Krashes et al., 2011), thus we used home cage food intake as a functional readout of successful expression of hM3Dq DREADDs or soma-targeted channelrhodopsin (soCoChR2) expression in AgRP neurons. For hM3Dq DREADDs experiments, mice were administered CNO (1mg/kg) before food access was reinstated 1 hour later. Food and body weight were then recorded at 2 hr- and 5 hr-post re-feeding. Mice that showed an elevated food intake response at 2 hours were deemed eligible for inclusion in the experiment. For optogenetic experiments, we used the food intake values reported during the 30 minutes of stimulation given during the training period to prevent any pretraining conditioning effects of transferring mice to the testing room. This time frame (10 minutes home-cage pre-simulation + 20 minutes within-context stimulation) was sufficient to drive a significant increase in feeding behaviour in *Agrp*^cre/wt^ but not *Agrp*^wt/wt^ mice. Mice that repeatedly failed to eat in response to stimulation across the 3 sessions were excluded from the CIF experiment.

### Immunohistochemistry

Successful viral transfection was confirmed by the presence of reporter expression in the Arc of *Agrp*^cre/wt^ and the absence from *Agrp*^wt/wt^ (mcherry: AAV5-hSyn-DIO-hM4D(Gi)-mCherry and AAV5-hSyn-DIO-hM3D(Gq)-mCherry; or GFP: AAV9 hSynapsin-FLEX-soCoChR-GFP), which was detected by immunohistochemistry. Following behavioural experiments, mice were transcardially perfused with 0.05M phosphate buffered saline (PBS) followed by 4% paraformaldehyde (PFA) in PBS. Following overnight post-fixation in PFA, brains were transferred to a 30% sucrose in 0.1M PB solution for a minimum of 24 hrs prior to cryosectioning. Sections were cut on a cryostat at 35 µm and stored in cryoprotectant at -20^°^C prior to processing. Free-floating sections were washed in 0.1M phosphate buffer (PB) followed by blocking in 4% normal horse serum (NHS) in 0.1M PB + 0.3% Triton-X. Primary antibodies were added to the blocking solution (Rabbit anti-DsRed 1:1000, stock #632496, Takara Bio, Clonetech; chicken anti-GFP 1:1000, ab13970, Abcam) and incubated overnight at 4^°^C. The next day, sections were washed in 0.1M PB before secondary antibody incubation for 2 hr at room temperature (Alexafluor Goat anti-rabbit 594 or Goat anti-chicken 488; 1:400 in 0.1M PB). Sections were mounted onto a slide and coverslipped using Vectorshield antifade mounting medium with DAPI (Vector Laboratories, Newark, CA USA).

### Context-induced feeding (CIF) assay

The CIF assay is a rapid, validated task in which food consumption is linked by association with an environmental context. Once paired, this association drives food consumption in sated mice. It provides a framework for studying the underlying neurobiology of feeding in the absence of current hunger (Stern et al., 2020).

#### Habituation

Prior to training, age-matched mice were habituated to palatable Froot Loops® breakfast cereal (168 kJ/g; 84.3 % carbohydrate, 5.4 % protein, 3.6 % fat, Kellogg’s, Australia) in their home cage on 3 separate occasions at different times of day. On the day prior to training, mice were also habituated to 2 separate contexts for 20 minutes each, in the absence of any food. Order of exposure to the contexts was counterbalanced across treatment groups. The contexts were designed to be distinct sensory experiences, differing in location (room); colour, shape, and texture of the testing arena (round yellow washing tub with a smooth floor surface, versus a comparably sized grey rectangular storage crate with an inserted ribbed flooring); smell (one of the following essences - rosewater, vanilla, rum, or almond); as well as placement and type of dish used as a food receptacle (ceramic dish vs 50ml falcon tube lid or petri dish). All animals were habituated to handling and received a minimum of 3 daily i.p. injections prior to experimentation, or 3 days habituation to tethered cables for optogenetic studies.

#### Training/acquisition of context-feeding

Training sessions involved placing mice into one of the contexts (assigned context A) for 30 minutes each, repeated across 3 days. Training in context A was paired with roughly 600-800 mg of a chopped mixture of Froot Loops placed inside the food dish. All contexts were wiped down with warm water and 50ul of appropriate olfactory cue reintroduced for each training session. All mice were trained within a 4-hour window in the mid-light phase, with food removed 1 hour prior to training (to eliminate the effects of recent feeding bouts) and returned 1 hour after they were returned to their home cage. This was repeated across 3 consecutive days for all experiments.

#### Testing/recall of context-induced feeding

Mice were re-fed in their home cages and tested across 2 days, 24-48 hr following the final training session. The presence of a conditioned feeding response was evaluated by comparing 20-minute palatable food intake in context A (the training context) to 20-minute intake in context B (a familiar, but untrained context). Order of context exposure was counterbalanced across days, and tests in each context were performed at the same time of day to prevent any time-of-day variation in feeding. All testing was performed in the absence of any treatment to assess the effect of treatment on acquisition, with the exception AgRP^HM4Di^, which were tested at the recall stage and therefore received no CNO during training but only prior to recall.

### CIF with fed, fasted, and ghrelin-treated C57Bl/6J mice

Experiments were performed as outlined above with the following exceptions. C57Bl/6J mice were randomly assigned to the following groups for training: Fasted + saline (FASTED; n = 9), Fed + saline (FED; n = 10), and Fed + 1 mg/kg ghrelin (GHRELIN; n = 10). Prior to each of the training days, mice allocated to the FASTED group had no food access starting from 2 hr before light onset, for a total of 10-12 hr before each training day. Following training, mice were re-fed 1 hr following training. For the FED- and GHRELIN-treated groups, food was removed concurrent with i.p. injection, which was performed 30 min prior to training. Access to home-cage food was returned to all mice 1 hr after the training session.

### CIF in fasted mice with Gi DREADD inhibition

Experiments were performed as outlined above with the following exceptions. AgRP^WT^ and AgRP^HM4Di^ mice were fasted prior to each training day, as detailed above. Training was performed *in the absence of food* to evaluate the role of AgRP neurons in mediating conditioned feeding independent of food reinforcement. In the first experiment, 1 hour prior to training, all mice received an injection of CNO (3mg/kg in saline), with food removed at the same time. In the second experiment designed to test the role of AgRP neurons in CIF recall, the same mice were habituated to a novel, comparably palatable food (Kellogg’s® Nutri Grain® breakfast cereal; 163kJ/g; 65.2% Carbohydrate, 21.8% protein, 3.0% fat; Kellogg’s, Australia) in their home cages and exposed to a completely novel set of contexts prior to training. Training was otherwise identical, except no treatment was given. ***Test:*** all mice were tested for 20 minutes in each context under fed conditions, with the same food present in context A or B. In the second experiment to test recall, mice received CNO (3mg/kg) 1 hour prior to each of the testing days.

### CIF in fed mice with Gq DREADD activation

Experiments were performed as outlined above with the following changes. all mice were trained under *ad libitum* (fed) conditions during the mid-light phase. Food was removed and all mice received i.p. injections of CNO (1mg/kg) 1 hour prior to being placed into the context. All mice had access to Froot Loops in the training context.

### CIF in fed mice with optogenetic stimulation

#### Experiment 1 – Stimulation with food

Experiments were performed as outlined above with the following exceptions. Fed mice were tethered to the fiber optic cables and allowed to settle for 10 minutes in their home cage prior to transfer into the training context. Training sessions were 30 minutes in duration, which included an initial 20 minutes of photostimulation (20Hz, 1s ON, 3s OFF) followed by 10 minutes in the absence of photostimulation. This was intended to be enough time to drive an acute feeding response in fed mice, but additionally allow a period without photostimulation following food consumption. Testing was performed without stimulation.

#### Experiment 2 – Stimulation without food

Experiments were performed as outlined above with the following exceptions. Fed mice were transferred to the training room and received 10 minutes of pre-stimulation in the home cage (20Hz, 1s ON, 3s OFF) before transfer into the training context. Photostimulation was terminated 10 minutes into the 30-minute training session, which was performed in the absence of any food in the training context. The withdrawal of AgRP stimulation within the training context was designed to mirror the silencing of AgRP neurons that naturally occurs in response to food presentation, to assess whether conditioning could be achieved in the absence of food or post-ingestive feedback during training. The presence of CIF was evaluated in the absence of photostimulation.

### Video analysis of behaviour

CIF experiments were recorded for video analysis using a fixed overhead camera (Sony HDR-50 Action Cam). Videos were scored using Ethovision tracking software (Noldus, US) to determine frequency, duration (s) and latency (s) for each pre-defined zone, as well as total session activity expressed as velocity (cm/s) and distance travelled (cm). For the CIF experiments, the zones included the food zone (immediately surrounding the food receptacle), as well as a ‘food approach’ zone extending an approx. 5 cm radius outside the food zone, and the remaining floor zone. Where described, behavioural barcoding across the training and test was performed on the videos using a combination of manual scoring and an in-house python analysis script, to establish the breakdown in behaviours in each zone over time.

### Statistical Information

Data were analysed using Prism 9 for MacOS (GraphPad Software, USA). Comparisons involving 2 factors (e.g. context, genotype) were analysed by 2-way ANOVA and Tukey’s post hoc test for multiple comparisons. Experiments involving within-animal comparisons were analysed using 2-way RM ANOVA followed by Sidak’s post hoc test for multiple comparisons. Tests comparing genotype alone for a single time point were performed using an unpaired t-test. Significance for all tests was reported at the level of p < 0.05.

## Results

### Mice trained under fasted, but not fed or ghrelin-treated conditions acquire context-induced feeding

To confirm previous reports that a fasted state during training sessions was necessary for mice to acquire CIF (Stern et al., 2020), we optimised an adapted protocol to accommodate potential variability in daily treatments. Further, we extended these findings by including a ghrelin-treated group to control for the effect of acute food intake during training, given both ghrelin and fasting are known to act on AgRP neurons to drive feeding (**Figure 1A**). Compared to Fed controls, Fasted mice ate significantly greater amounts across each of the training sessions (p < 0.05; **figure 1B**). While Ghrelin mice ate more than Fed mice on average across the 3 days (unpaired t-test; p < 0.05), this was not significant in the overall comparison, with a trend toward increased intake evident only on the first day (p = 0.073). When all mice were re-fed and later tested for CIF, only the Fasted group displayed a CIF response based on food intake in context A vs context B (p = 0.007; **figure 1C**) or when context discrimination was expressed as the individual difference in food intake between context A vs context B (p = 0.034; **figure 1D**). Fasted mice also showed the strongest relationship between average training food intake and context discrimination for the CIF test (R^2^ = 0.348), although this was not significant (A-B; **figure 1E**).

**Figure 1.**
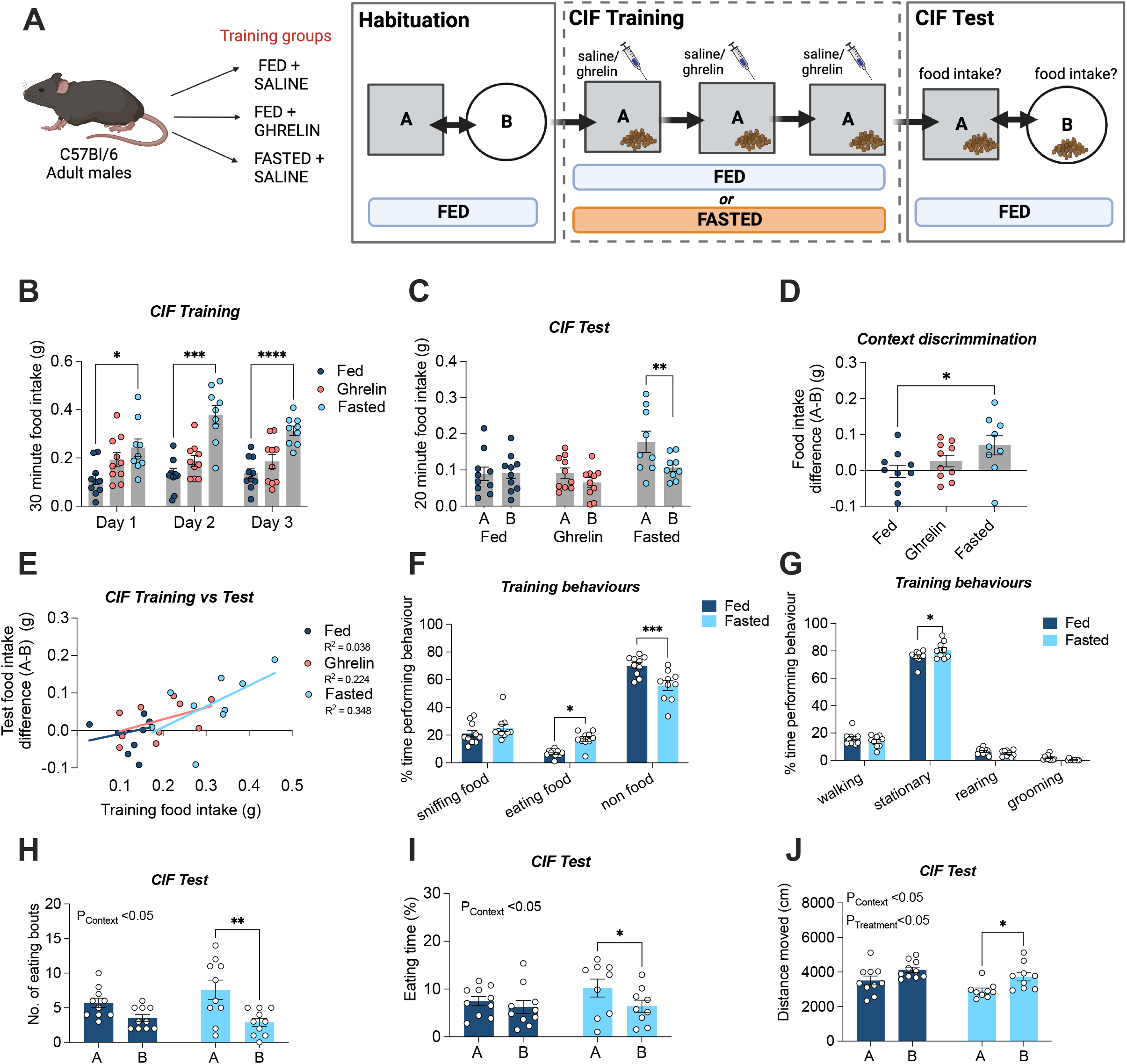
Context induced feeding (CIF) in C57Bl/6J mice trained under fed, ghrelin or fasted conditions. (A) Adult C57Bl/6J mice were split into 3 groups for CIF training: fed + saline (i.p) (n = 10), fed + ghrelin (1mg/kg, i.p) (n = 10), and fasted + saline (i.p) (n = 9). Following habituation to contexts A and B, all mice received 3 x days of training in context A paired with palatable froot loops and were then tested for a CIF response by comparing intake in context A to B; created with BioRender.com. (B) CIF training food intake for fed, ghrelin and fasted groups across each of the 30-minute training sessions. (C) The presence of a CIF response was evaluated by comparing food intake in context A to B during 20-minute test sessions. (D) CIF test data expressed as the difference in food intake in context A and B (A-B; one-way ANOVA with Dunnett’s post hoc test for multiple comparisons). (E) Relationship between food consumed during training (x axis) and context discrimination at test. (F) Breakdown of % time engaging in food- and non-food related behaviours during training in fed and fasted groups. (G) Breakdown of % time engaging in specific non-food behaviours during training (walking, stationary, rearing, grooming) in fed and fasted groups. (H) Number of eating bouts during CIF testing in A and B in fed and fasted groups. (I) % time eating during CIF testing in A and B in fed and fasted groups. (J) Locomotor activity (total distance travelled) during testing in context A and B for fed and fasted groups. All values are expressed as mean +/- SEM. Data analysed by 2-way RM ANOVA (main effects stated on graphs) with Sidak’s post hoc test for multiple comparisons, unless otherwise stated. *p < 0.05, **p < 0.01, ***p < 0.001, ****p < 0.0001.

Given there were no differences between Fed and Ghrelin groups in acquiring CIF, we profiled the behaviours of mice from Fed and Fasted groups on the first and last training days (Day 1 and Day 3) and the two test days (Contexts A and B), focusing on the first 10 min of each trial when changes in appetitive behaviour are likely to be most prominent. As expected, Fasted mice exhibited the greatest food-oriented behaviour during training; spending significantly more time eating food (p = 0.01; **figure 1F**), and an overall reduction in time engaging in non-food related behaviours (p value; **figure 1F**). When we separately characterised the non-food related behaviours, both groups spent most of their time stationary, but this was significantly higher in the Fasted group due to the fact they spent more time consuming food (p = 0.030; **figure 1G**). There were no differences in time spent sniffing food (**figure 1F**), walking, rearing, or grooming (all p’s > 0.05; **figures 1G)**.

When food intake in context A was compared to B in the CIF test, mice trained in the Fasted group engaged in more bouts of feeding (0.001; **figure 1H**) and spent significantly more time eating (p = 0.040; **Figure 1I**), whereas mice trained in the Fed group displayed similar feeding behaviour in both contexts (p’s > 0.05). Overall mice displayed greater locomotor activity in context B compared to A (p = 0.005), but a significant difference between contexts was only apparent in the Fasted group (p = 0.041; **figure 1J**). Altogether, these results are consistent with Fasted mice engaging in more food-oriented behaviours during training, which led to the formation of food-specific conditioned feeding behaviour at test.

### AgRP neurons are necessary to acquire CIF in fasted mice

Both fasting (negative valence) and consumption of palatable food (positive valence) in a context during training influence the acquisition of CIF (Stern et al., 2020). Therefore, we asked whether AgRP neurons are necessary to acquire CIF in fasted mice, independent of food consumed during training. To do this, we chemogenetically suppressed (**figure 2A and 2B**) AgRP neuronal activity during CIF training in the absence of food and later tested for a CIF response under fed conditions (**figure 2C**). AgRP^WT^ controls showed intact CIF (p = 0.011) exhibiting significantly greater food intake in context A compared to B. In contrast, AgRP^hM4Di^ mice (tested in the absence of CNO) showed no preference for either context (p = 0.7855; **figure 2D**); highlighting a necessary role for AgRP neurons in acquiring CIF, independent of palatable food intake.

**Figure 2.**
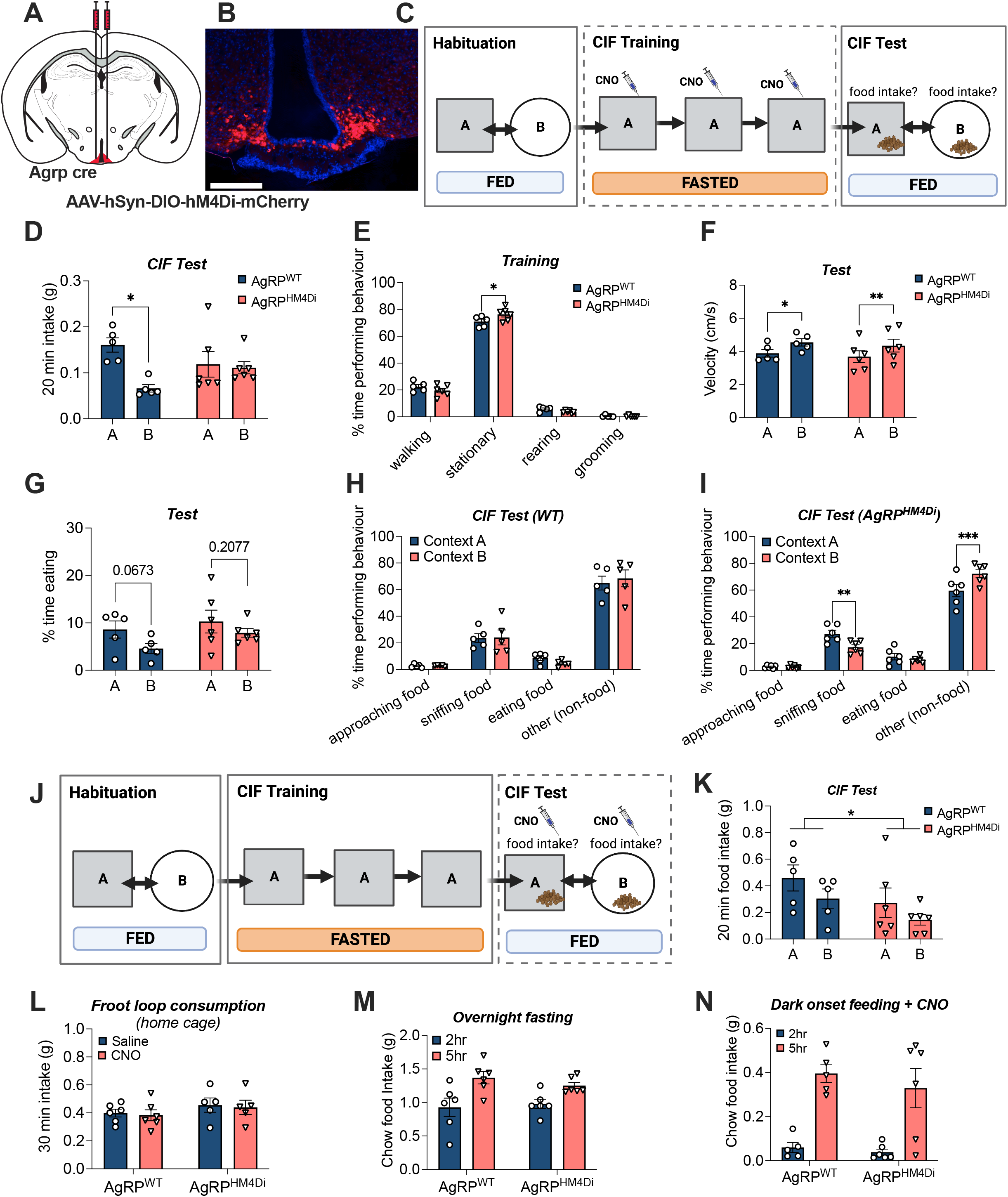
Chemogenetic inhibition of AgRP neurons in mice fasted during training. (A) Schematic of injection site for AAV-hsyn-DIO-hM4D (Gi) DREADDs into the Arc of AgRP-ires-cre (n = 6) and AgRP-WT littermates (n = 5), creating AgRP^Gi^ and AgRP^WT^; respectively. (B) Representive image showing expression of hM4Di DREADDs in the Arcuate nucleus (Arc) of AgRP-ires-cre mice; sacle bar 200 µm. (C) Experimental overview for AgRP inhibition in fasted mice during training created with BioRender.com. Mice were habituated to contexts A and B and then trained under fasted conditions in the presence of CNO (3mg/kg) for 3 days. The presence of a context-induced feeding (CIF) response was evaluated under fed conditions by comparing intake of palatable froot loop intake in A to B. (D) CIF without CNO; froot loop intake in A and B during 20-minute tests in each context. (E) Behavioural characterisation of behaviours during training, in the absence of food. (F) Locomotor activity expressed as velocity (cm/s) in A and B at test. (G) Proportion of time spent eating (%) in contexts A and B at test. (H) Breakdown of time engaging in food- and non-food-related behaviours for AgRP^WT^ mice, and (I) AgRP^Gi^ mice. (J) Experimental overview for AgRP inhibition in fed mice at test created with BioRender.com. (K) CIF with CNO; froot loop intake during 20-minute tests in each context. (L) 30-minute froot loop consumption in the home cage, in the presence of CNO or saline (counterbalanced). (M) Food intake at 2hrs and 5hrs following CNO pre-treatment prior in overnight fasted mice. (N) Food intake following CNO treatment in the late light phase, prior to regular feeding. Values are presented as mean +/- SEM. *p < 0.05, **p<0.01, trends presented as actual p values.

With respect to the behavioural differences underlying this effect, during the first 10 min of training there were no differences between AgRP^WT^ and AgRP^HM4Di^ mice in the proportion of time spent walking, rearing, or grooming. However, chemogenetic inhibition of AgRP neurons led to a significant increase in the time spent stationary (p = 0.029; **figure 2E**), which could not be attributed to food consumption. Under test conditions, both groups expressed increased locomotor activity (velocity; cm/s) in context B, compared to context A (p = 0.0004; **figure 2F**), consistent with a novelty-induced locomotor effect. Previously fasted AgRP^WT^ mice had a trend towards reduced time eating in context B (p = 0.067), but otherwise displayed no difference in food- or non-food related behaviours at test (all p’s > 0.05; **figures 2G & 2H**). On the other hand, AgRP^hM4Di^ mice spent similar time eating in contexts A and B (**figure 2G**), but spent significantly greater time sniffing food in context A (A vs B; p = 0.0046; **figure 2I**) and overall spent less time in non-food related behaviours in context A compared to B (p = 0.0005; **figure 2I**). Altogether, these data suggest that AgRP neurons contribute to the fasting-mediated CIF effect independent of food intake during training.

We also assessed whether AgRP neurons were necessary for expression of CIF at test by suppressing AgRP activity with CNO prior to the CIF test (**figure 2J**). Although in this experiment there was an overall effect of increased feeding in context A (p = 0.0483), neither AgRP^WT^ mice nor AgRP^hM4Di^ groups significantly discriminated between context A and B at test (**figure 2K**), suggesting AgRP neurons play a greater role in the acquisition, rather than recall of CIF.

Given that no Froot Loops were offered during training, we additionally assessed the role of CNO-driven inhibition on acute consumption of Froot Loops in the home cage under matching fasted conditions. There were no differences in consumption between AgRP^WT^ and AgRP^HM4Di^ mice in response to either saline or CNO (**figure 2L**), suggesting suppression of AgRP activity does not influence unconditioned feeding of the palatable food in a home cage environment. There were also no differences in home cage chow intake when CNO was administered prior to re-feeding after an overnight fast, at 2- or 5-hrs (**figure 2M**), nor were there differences in feeding responses to CNO under another condition when AgRP activity is typically high – at the onset of the dark phase (**figure 2N**), suggesting the effect observed with CIF is separate from the acute effects of palatable and regular chow intake in the home cage.

### Acute temporal optogenetic control of AgRP neurons drives CIF

Both chemo- and opto-genetic activation of AgRP neurons drive acute feeding in fed mice (Aponte et al., 2011; Krashes et al., 2011). Therefore, we next set out to understand whether activation of AgRP neurons during training was sufficient for fed mice to acquire CIF, in the presence of food. Firstly, AgRP-IRES-cre mice expressing excitatory DREADDs (AgRP^hM3Dq^) (**figure 3A and 3B**) and their wild-type littermates (AgRP^WT^) underwent CIF under fed conditions, with CNO administered daily prior to training sessions (**figure 3C**). During training, AgRP^hM3Dq^ mice ate significantly greater amounts in response to CNO (average training intake p < 0.0001), indicating CNO treatment was successfully targeting AgRP neurons to drive a feeding response (**figure 3D**). However, when tested for CIF in the absence of CNO, AgRP^hM3Dq^ mice did not eat more than controls, nor did they show a CIF response (**figure 3E**). Moreover, these mice showed a negative relationship between acute and conditioned feeding responses (training versus test food intake in context A), in contrast to AgRP^WT^ mice whose training and test intake showed an expected positive relationship (p < 0.0001; **figure 3F**). Therefore, although chemogenetic stimulation of AgRP neurons was sufficient for acute feeding within-context, this did not translate to a context-conditioned feeding response at test. The acute feeding response in fed mice was also confirmed in a home cage environment with standard chow, where AgRP^hM3Dq^ mice displayed a pronounced feeding response to CNO (p < 0.0001) in the mid-light phase (**figure 3G**).

**Figure 3.**
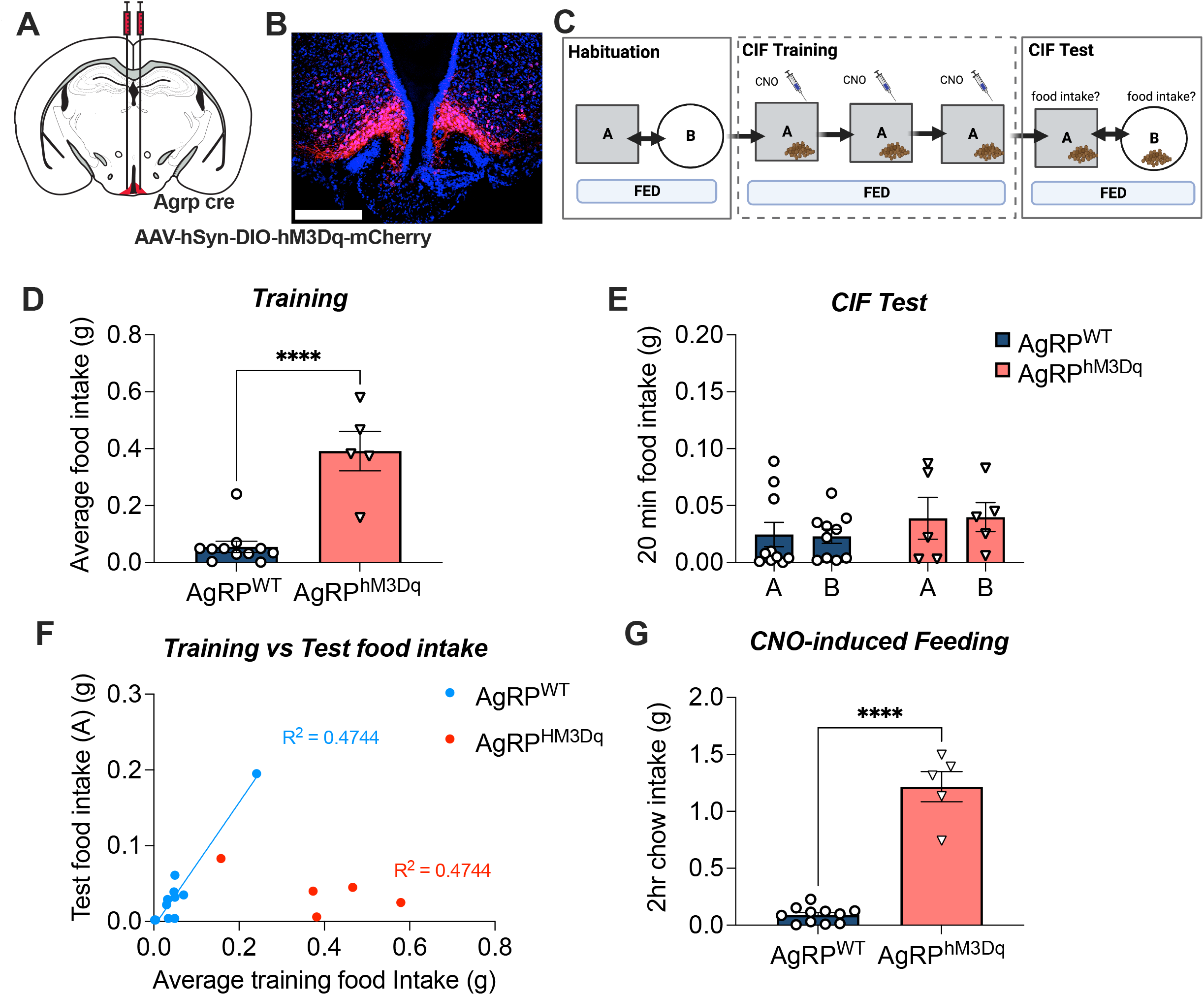
Chemogenetic stimulation of AgRP neurons in fed mice during training. (A) AgRP-ires-cre and AgRP-WT littermates received bilateral injections of AAV5-hsyn-DIO-hM3Dq-mcherry into the arcuate nucleus creating AgRP^HM3Dq^ (n = 5) and AgRP^WT^ (n = 11); respectively. (B) Representative image of AAV-hsyn-DIO-hM3Dq-mCherry in AgRP-ires-cre mice. (C) Experimental overview created with BioRender.com. All mice were trained under fed conditions and received injections of CNO (1mg/kg) 1hour prior to training sessions. The presence of a context-induced feeding (CIF) response was evaluated under fed conditions by comparing intake of froot loops in contexts A and B. (D) Average froot loop intake (g) across each of the 30-minute training sessions. (E) Context-induced feeding test; 20-minute froot loop intake in contexts A and B under fed conditions. (F) Relationship between training food intake and test food intake in context A. (G) Representative image of injection site placement (mcherry reporter expression) in AgRP-ires-cre mice. Values are presented as mean +/- SEM, ****p < 0.0001.

Because food and food cues rapidly inhibit AgRP neurons (Betley et al., 2015; Chen et al., 2015; Reichenbach et al., 2022; Su et al., 2017), we questioned whether the sustained chemogenetic activation of AgRP activity, coupled with the lack of temporal AgRP activity changes prevented the acquisition of CIF, even though activation increased food intake during training. Therefore, we repeated the CIF assay in fed mice expressing AgRP-dependent soma-targeted channelrhodopsin (AgRP^SoCoChR^) or controls (AgRP^WT^), to concentrate ChR2 on AgRP cell bodies in the ARC. We used a short period of photostimulation for 20 min within-context for each 30-minute training session (**figure 4A and 4B**). Photostimulation significantly elevated feeding in AgRP^SoCoChR^ mice during training (p = 0.018; **figure 4C**) and when assessed for CIF at the test stage without photostimulation, AgRP^SocoChR^ mice also displayed a CIF response (p = 0.028) that was absent in AgRP^WT^ mice (A vs B; p = 0.574; **figure 4D**). This suggests the short period of within-context AgRP photostimulation was sufficient to acquire CIF. Although both AgRP^hM43Dq^ mice (f**igure 3)** and AgRP^SocoChR^ mice increased feeding responses during training, only the temporally-specific photostimulation and cessation permitted development of a conditioned feeding response to context. This is highlighted by the positive relationship between training food intake and test food intake in context A in AgRP^SocoChR^ mice (**figure 4E**) but not AgRP^hM43Dq^ mice (see **figure 3F**).

**Figure 4.**
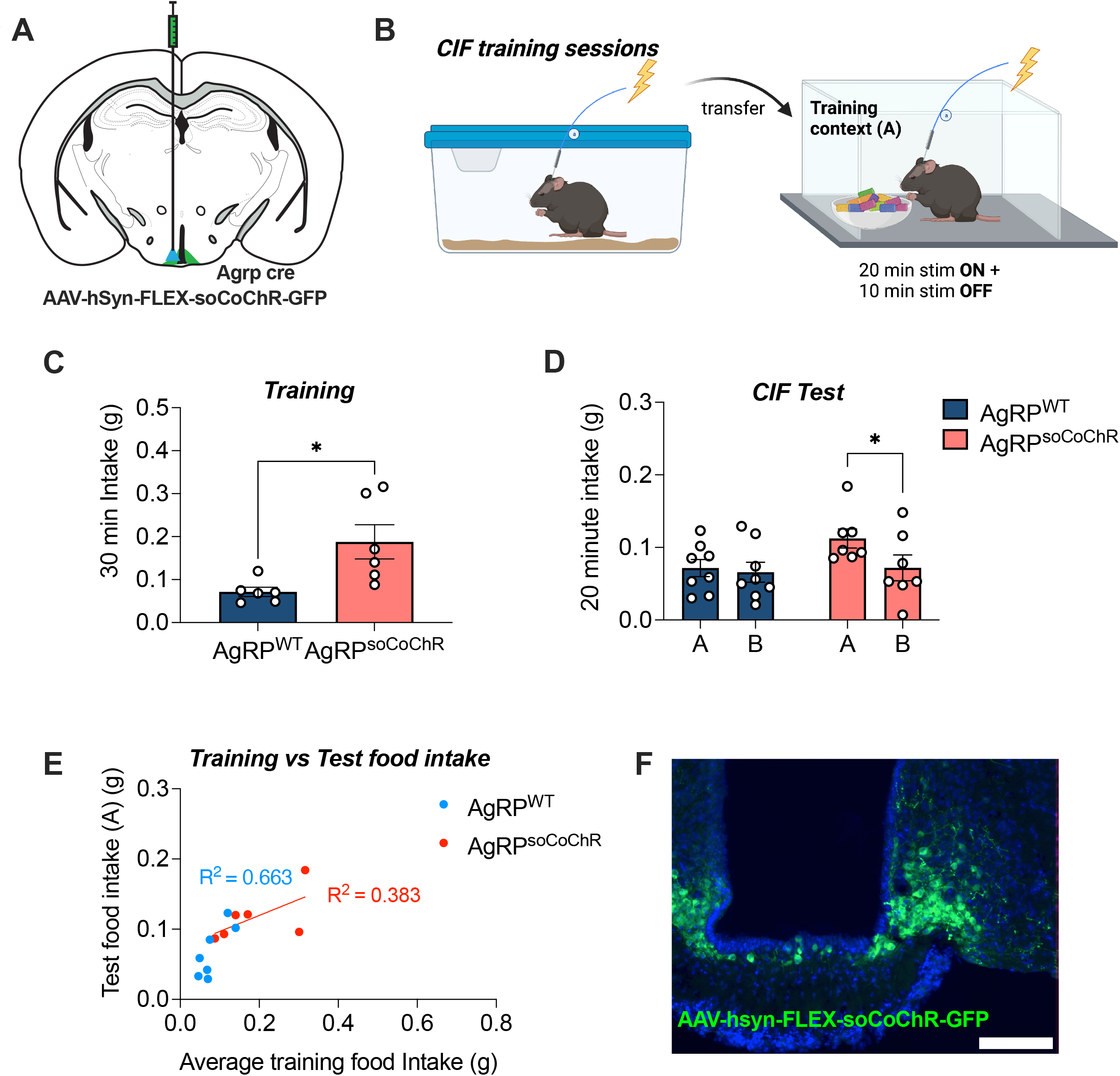
Within-context optogenetic stimulation of AgRP neurons during training. (A) AgRP-ires-cre and AgRP-WT littermates received bilateral injections of AAV9-hsyn-FLEX-soCoChR-GFP into the arcuate nucleus creating AgRP^soCoChR^ (n = 6) and AgRP^WT^ (n = 6); respectively. An indwelling fiber optic cannula was implanted unilaterally above the injection site. (B) Experimental overview created with BioRender.com. Mice were habituated to contexts A and B and then trained under fed conditions. 20Hz Photostimulation (1s on, 3s off) was delivered for the first 20 minutes of the 30-minute training session, with palatable froot loops available. The presence of a context-induced feeding (CIF) response was evaluated under fed conditions by comparing intake of froot loops in contexts A and B. (C) Froot loop intake averaged across each of the 30-minute training sessions. (D) Context-induced feeding test; 20-minute froot loop intake in contexts A and B under fed conditions. (E) Relationship between training food intake and test food intake in context A. (F) Representative image of GFP reporter expression in AgRP-ires-cre mice. Values are presented as mean +/- SEM, *p < 0.05.

### Withdrawal of artificial AgRP stimulation in the absence of food is sufficient for CIF

The results of the previous experiment suggested the timing of AgRP neuron activation relative to context exposure is critical for conditioned, but not normal feeding responses in fed mice. Recent data shows that AgRP neurons are rapidly inhibited by food-predictive cues, with the greatest response to calorie-rich food in hungry mice. These data are in line with the idea that temporally-defined inhibition of AgRP neuronal activity upon food-cue presentation influences learning (Berrios et al., 2021; Reed et al., 2022; Su et al., 2017). Under physiological conditions, post-ingestive feedback from calories maintains the suppression of AgRP neural activity; however, the role of the acute silencing to food-predictive cues independent of post-ingestive feedback remains unclear.

In the final set of experiments, we set out to artificially recreate this pattern of AgRP activity (**figure 5A**) in the absence of hunger or post-ingestive feedback from food consumption, to determine whether activity dynamics alone would be sufficient for mice to acquire CIF. Fed AgRP^SocoChR^ and AgRP^WT^ mice received daily training sessions in context A with a brief period (10 minutes) of home-cage photostimulation (pre-stim), that continued for the first 10 minutes in the training context. Termination of photostimulation within context A mimicked the suppression of AgRP activity typically caused by the presentation of food and food-predictive cues. Remarkably, the termination of AgRP photostimulation within context A – even in the absence of food during training - was sufficient to drive CIF in AgRP^SocoChR^ mice, when palatable food was later offered in both contexts at test (A vs B; p = 0.005). Notably, this effect that was absent in AgRP^WT^ mice (A vs B; p = 0.619) (**figure 5B**). Because no food was offered during training, we measured home cage chow intake following training sessions, but this was unaffected compared to controls at 24- or 48-hours (p > 0.05; **figure 5C**). In contrast, AgRP photomstimulation in AgRP^SocoChR^ mice increased home cage chow food intake compared to AgRP^WT^ mice control mice (**figure 5D**). Thus, the removal of artificially-driven AgRP activity inside the context and in the absence of fasting or the presentation of food, is sufficient for the later expression of context-induced feeding. These data highlight that hunger-driven learning requires an acute and maintained suppression (at least 20 minutes in context) of AgRP activity in response to sensory cues that predict food availability and consumption.

**Figure 5.**
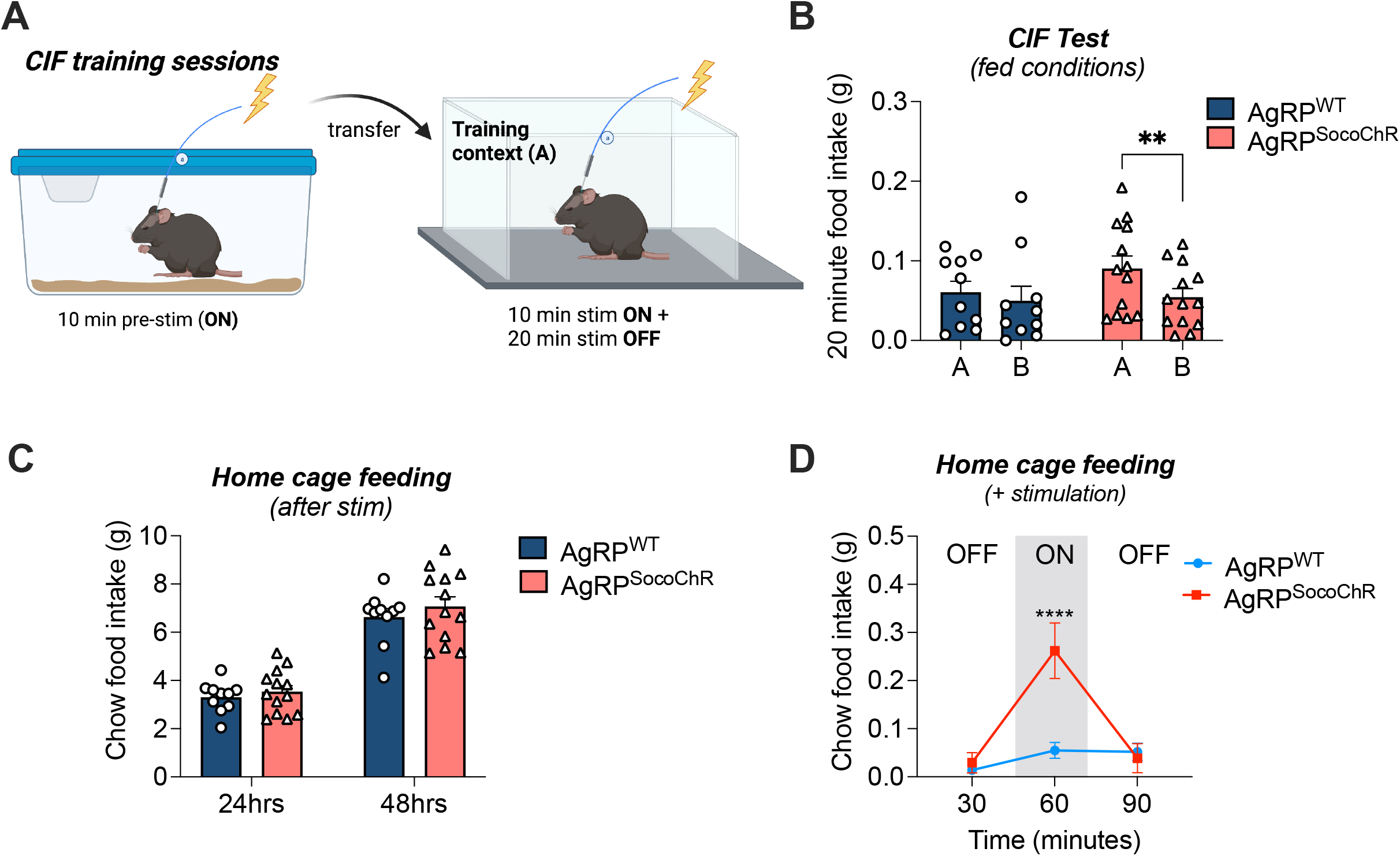
Withdrawal of optogenetic stimulation of AgRP neurons in the training context. AgRP-ires-cre and AgRP-WT littermates received bilateral injections of AAV9-hsyn-FLEX-soCoChR-GFP into the arcuate nucleus with an indwelling fiber optic cannula implanted above the injection site, creating AgRP^WT^ (n = 10) and AgRP^SocoChR^ (n = 13) groups, respectively. (A) Schematic created with BioRender.com outlining conditions of optogenetic stimulation given on each of the training days for the context-induced feeding experiment. Fed mice received 10 minutes of pre-stimulation in the home cage, followed by 10 minutes stimulation in the training context and an additional period without stimulation. Training was performed in the absence of food. (B) Context-induced feeding Test. Froot loop intake in 20-minute testing sessions in contexts A and B. (C) Home cage feeding during the training period, measured at 24- and 48-hrs following the first training session. (D) Home cage chow food intake measured at 30-minute intervals; prior to photostimulation (OFF), 30 minutes post 20Hz stimulation (ON), and 30 minutes after stimulation offset (OFF). Data analysed with 2-way RM ANOVA, and Sidak’s post hoc test for multiple comparisons. Values are presented as mean +/- SEM, **p < 0.01, ****p < 0.0001.

## Discussion

In this series of studies, we used optogenetic and chemogenetic manipulations of AgRP neurons to examine the reciprocal interactions between hunger and memory in a context-induced feeding (CIF) assay. We first demonstrated fasting was necessary for successful acquisition of CIF despite offering the same palatable food in an alternative context at test, and fed controls receiving an otherwise identical treatment. Our results are consistent with previous studies in mice and rats showing a similar prerequisite of food restriction for learning (Petrovich et al., 2007; Stern et al., 2020; Weingarten, 1983). Having established a working model to examine CIF, we sought to understand a neural mechanism through which fasting facilitates a context-conditioned feeding response. We hypothesised that discrepancies between fed and fasted mice could be attributed to the activity of AgRP neurons, which are more active under fasted conditions (Mandelblat-Cerf et al., 2015) and are silenced by food and food-predictive sensory cues (Betley et al., 2015; Mandelblat-Cerf et al., 2015; Reichenbach et al., 2022; Su et al., 2017). Indeed, population recordings of AgRP neurons show that palatable food has a greater effect on AgRP neuron silencing in fasted compared to fed mice (Chen et al., 2015), suggesting the magnitude of AgRP suppression in response to food and/or food cues is critical to acquire CIF. This is further reinforced by the lack of CIF in fed mice.

Further, we extended these findings to show that i.p. administration of ghrelin is not sufficient to acquire CIF under these conditions, despite ghrelin having known activity on AgRP neurons (Andrews et al., 2008; Beutler et al., 2020; Lockie et al., 2018; Reichenbach et al., 2022). A further finding is that palatable food consumption during training was not necessary for the fasting-mediated effect, because fasting alone could similarly precipitate CIF at test, as previously demonstrated (Stern et al., 2020). That fasting alone is sufficient for acquisition of CIF, which depends on AgRP activity as shown by chemogenetic inhibition, indicates that AgRP neurons contribute to conditioned hunger, an appetitive drive that potentiates feeding when food is later offered under fed conditions. Collectively, this highlights a critical role for AgRP *neural firing*, as opposed to increased consumption following AgRP activation, in acquiring a context-induced feeding memory. This observation is further underscored by the inability of chemogenetic AgRP activation to drive a CIF, despite feeding responses within the training context, as previously described (Aponte et al., 2011; Krashes et al., 2011). Intriguingly, our results demonstrated that only the temporally controlled optogenetic conditions were appropriate to acquire a context-dependent memory. In support of this, we showed that artificially mirroring the temporal dynamics of AgRP activity in fed mice is sufficient to acquire CIF *in the absence of food*, suggesting a transient shift of AgRP neurons from the active to inactive state in the training context drives fasting-mediated CIF.

One notable observation was that sustained activity of AgRP neurons using hM3DGq DREADDs did not facilitate acquisition of context-induced feeding, and it is possible that a sustained increase of AgRP activity mediated by ghrelin administration may have had the same effect. Indeed, another study utilising a slightly modified version of conditioned feeding, showed that activation of AgRP neurons with hM3DGq DREADDs in fed mice increased the feeding response within context, but failed to produce a subsequent context-induced feeding response (Mohammad et al., 2021). The most important feature of chemogenetic approaches is the relatively long 6-hour half-life after CNO injection (Roth, 2016). In comparison to normal fasted situations in which AgRP neurons are highly active and rapidly inhibited in response to food, chemogenetic activation of AgRP neurons in fed mice causes a prolonged activation despite food presentation and food cues. Thus, we suggest this chemogenetic approach creates a temporal mismatch between persistently high AgRP neuronal activity and food consumption which does not replicate physiological changes in AgRP firing properties. However, an optogenetic approach can maintain the required firing properties of AgRP neurons in vivo, and in our studies photostimulation of AgRP neurons limited to 20 minutes (out of 30 mins) in the training context was sufficient to induce CIF in fed mice, suggesting that a fall in activity (from photostimulation ON-OFF) was necessary to learn the task. These studies demonstrate an important distinction between approaches to study the function of AgRP neurons, and suggest caution should be applied when interpreting and comparing data from AgRP DREADD and optogenetic studies.

The rapid silencing of AgRP neurons naturally occurs in response to the presentation of food, or food-predictive cues such as the sight or smell of a known food source (Betley et al., 2015; Chen et al., 2015; Su et al., 2017). The rapid nature of this inhibitory activity suggests afferent neural circuits, as opposed to gut feedback, are responsible although post-ingestive gut feedback is required for sustained inhibition of AgRP neurons. Chen and Knight proposed several possible explanations for an acute silencing signal, including to connect current experiences with future outcomes (Chen and Knight, 2016). Our data are consistent with this possibility. For example, the inhibition of AgRP neuronal activity with Gi DREADDs prevented the acquisition of fasting-induced CIF. Moreover, our temporally confined optogenetics suggest the timing of the ‘OFF’ signal for AgRP neurons, as would naturally occur with food presentation, acts to link the change in AgRP activity to the current associated contextual sensory cues. Our most striking finding was that sustained cessation of AgRP photostimulation, in the absence of food, was sufficient to acquire a CIF response. The key reason for this may relate to acute transient inhibition vs prolonged cessation of AgRP activity, as we postulate the sustained cessation of AgRP photostimulation mimics post-ingestive metabolic feedback, which confirms calorie consumption and resets AgRP firing to lower rates (Beutler et al., 2017; Goldstein et al., 2021; Su et al., 2017). The fact that CIF could be acquired through withdrawal of AgRP activity alone supports the idea of an acute silencing signal serving as a link between current and future outcomes. In this manner, the suppression of AgRP neural activity may be a novel neural representation of positive reinforcement, which aligns with the common use of food rewards in reinforcement learning paradigms. Moreover, the acute silencing of AgRP neurons might serve as a predictor in a prediction-error signal requiring post-ingestive feedback for evaluation and updating. However, there are various other stimuli that are known to silence AgRP activity, such as the cessation of exercise (Miletta et al., 2020), thermal pain (Alhadeff et al., 2018), pup reunion to dam (Zimmer et al., 2019), alcohol (Alhadeff et al., 2019) and cold temperature (Deem et al., 2020; Yang et al., 2021b). The acute silencing signal alone may have important consequences for learning that do not require post-ingestive feedback. Thus, inhibition of AgRP neurons to non-food cues may provide additional information about an environment, although future experiments are required to investigate this further. We also show here that chemogenetic inhibition of AgRP neurons reduced activity in the absence of food, resulting in increased time spent stationary. This result is in line with previous reports that AgRP neurons increase locomotor activity in the absence of food (Krashes et al., 2011) and consistent with the proposal that acute AgRP silencing prior to post-ingestive feedback promotes a state-transition from appetitive to consummatory behaviour (Chen and Knight, 2016).

It is interesting to note that AgRP neurons drive learning in passive conditioning studies, such as those presented here, and in other studies using operant conditioning (Betley et al., 2015; Chen et al., 2016; Krashes et al., 2011; Reichenbach et al., 2022), suggesting they facilitate appetitive conditioning more generally. This is consistent with the idea that hunger has been used as a key motivational tool for learning in behavioural neuroscience for decades and uncovers an important and largely unexplored role for AgRP neurons in learning and memory, as we recently discussed (Reed et al., 2022). Moreover, the fact that activation of AgRP neurons with Gq DREADDs drives food seeking and consumption but not CIF, whereas the temporal suppression of AgRP neurons induces CIF in the absence of food intake, highlights for the first time that AgRP neurons may promote motivation and learning through distinct downstream pathways. The nature of these pathways remains to be elucidated, however studies using CIF have shown elevated activity of the insular cortex, lateral hypothalamus, central amygdala and lateral septum following CIF (Stern et al., 2021). AgRP neurons influence the firing of insular cortex neurons to visual cues that predict food availability via a relay from the paraventricular thalamus and basolateral amygdala (Livneh et al., 2017), suggesting activation of AgRP neurons modulates the encoding of information across various brain regions. AgRP neuron activation also increases motivation by acting on dopaminergic pathways in the nucleus accumbens (Alhadeff et al., 2019; Reichenbach et al., 2022); a site which is also involved in associative reward memory (Day et al., 2007). Moreover, AgRP neurons are regulated by direct inputs from DMH GABA-ergic neurons co-expressing the leptin receptor (lepr) and prodynorphin (pdyn) (DMH^lepr/pdyn^), and likewise, this projection is active in response to food-predictive cues; suggesting DMH^lepr/pdyn^ neurons contribute to the early and rapid silencing of AgRP neurons that occurs in response to food cues (Garfield et al., 2016). Inhibition of these upstream pathways also increases the time taken to acquire a visual discrimination task for food rewards (Berrios et al., 2021) implying that AgRP neurons influence the acquisition of this task, although a specific role for AgRP neurons was not examined in this study.

In summary, our results show that AgRP neurons mediate the acquisition of CIF, and importantly, highlight that the timing of AgRP silencing relative to context exposure is likely responsible for CIF acquisition. This silencing of AgRP neurons may confer a learning signal in the absence of post-ingestive feedback, as evidenced by the presence of conditioned responses in those experiments performed in the absence of food. More broadly, our results highlight that dynamic changes in AgRP activity, rather than AgRP-mediated food consumption, influence learning and memory, and pinpoints a neural population responsible that the well-established effects of hunger on associative learning. We suggest that the magnitude of AgRP suppression to a salient signal, as well as the length of that suppression, underlies the learning potential embedded within the firing properties of AgRP neurons. Furthermore, finding a way to disrupt the formation of conditioned feeding memories may be useful to help reduce overeating, food cravings and the development of obesity.

## Acknowledgements

This study was supported by an NHMRC grant and fellowship to ZBA (1126724, 1154974) and NHMRC grant and fellowship to SHL (1147700, 1072364). We acknowledge that BioRender was used to produce elements incorporated in the figure and graphical abstract (Biorender.com).

